# A honey bee-associated virus remains infectious and quantifiable in postmortem hosts

**DOI:** 10.1101/2024.08.08.607215

**Authors:** Alexandria N. Payne, Vincent Prayugo, Adam G. Dolezal

**Author notes:** Author to whom correspondence should be addressed: Alexandria N. Payne.

## Abstract

Corpse-mediated transmission is a potentially viable route through which naïve hosts can become infected, but its likelihood for honey bee-associated viruses is largely unknown. While these viruses can be easily detected in deceased bees, it remains unclear if they stay infectious within postmortem hosts or if enough viral RNA degradation—and subsequently virus inactivation—occurs post-host death to render these viruses inviable. This knowledge gap has important implications for how researchers perform honey bee virus studies and for our general understanding of honey bee virus transmission. To better understand the resiliency of honey bee-associated viruses within deceased hosts, we first tested the hypothesis that postmortem specimens, stored in colony-normal temperature and humidity conditions, can be reliably used to quantify virus abundance. To determine this, we experimentally-infected adult honey bees with Israeli acute paralysis virus (IAPV) and then measured the virus levels of individuals sampled live or at different postmortem timepoints (4–, 12–, 24–, and 48–hours post-death) using RT-qPCR and a standard curve absolute quantification method. We found no significant differences based on when bees were sampled, indicating that postmortem honey bees are statistically comparable to using live-sampled bees and can be reliably used to quantify absolute IAPV abundance. We then performed a follow-up experiment that determined whether or not the IAPV detected in postmortem bees remained infectious over time. We found that IAPV extracted from postmortem bees remained highly infectious for at least 48–hours post-death, indicating that any viral RNA degradation that may have occurred during the postmortem interval did not adversely affect IAPV’s overall infectivity. The results from this study suggest that IAPV is more resilient to degradation than previously assumed, support the use of postmortem bees for downstream IAPV analyses, and indicate that postmortem hosts can act as sources of IAPV infection for susceptible individuals.

## 1. INTRODUCTION

The potential for postmortem hosts to remain sources of viral infection depends on the extent of virus degradation that occurs prior to transmission. This is influenced by factors such as environmental conditions and the structure of the virus itself (Fordyce et al., 2013). For honey bee (*Apis mellifera*)-associated viruses, of which most are non-enveloped, +ssRNA viruses (Brutscher et al., 2016; Chen and Siede, 2007; Grozinger and Flenniken, 2019), corpse-mediated transmission has the potential to occur, namely through corpse-removal behavior (López-Riquelme and Fanjul-Moles, 2013) performed by undertaker honey bees (Trumbo et al., 1997; Visscher, 1983) and by continuous accumulation of virus particles by the ectoparasitic mite *Varroa destructor* (Di Prisco et al., 2011; Posada-Florez et al., 2019; Traynor et al., 2020) when feeding on hosts that have died as a result of infection. However, whether these honey bee-associated viruses remain infectious within a deceased host, and to what extent, remains largely unknown, despite its importance in better understanding how these viruses persist and are transmitted between honey bee individuals, colonies, and other non-*Apis* species (Dolezal et al., 2016; Payne et al., 2020; Yañez et al., 2020, 2012).

In general, RNA viruses have historically been considered highly susceptible to degradation when outside of a viable host environment, which is commonly reflected in the way specimens are sampled across different host-pathogen systems (Bustin and Nolan, 2004; Fleige and Pfaffl, 2006; Gallego Romero et al., 2014). For example, preferential sampling of live specimens is based on evidence that the postmortem interval (*i*.*e*., the time that has elapsed since an individual’s death) can significantly impact RNA integrity (Durrenberger et al., 2010), leading to compromised accuracy and reliability of downstream analyses (Koppelkamm et al., 2011; Lee et al., 2005; Zhu et al., 2017). The concern that postmortem sampling negatively impacts RNA integrity is evident within the field of honey bee research, as the “gold-standard” sampling method for honey bee-associated viruses calls for the collection of live specimens flash-frozen in liquid nitrogen and immediately stored at -80 ºC (de Miranda et al., 2013; Evans et al., 2013). Alternative sampling methods that deviate from this standard are typically considered suboptimal, as they can cause increased degradation of total RNA within a specimen (Chen et al., 2007; Evans et al., 2013; Forsgren et al., 2017). However, if postmortem specimens can be reliably used for viral analyses, such as quantifying absolute virus abundance, they present a sampling alternative that could offer more consistent comparisons between infected individuals while not further depleting the sampling pool intended for other analyses (e.g., assessing survivorship).

When postmortem bees are used for viral analyses, they are often viewed as a contingency, with only positive results being deemed reliable due to concerns of false negatives caused by RNA degradation (de Miranda et al., 2013; Evans et al., 2013). In fact, these previous studies explicitly discourage the use of postmortem bees to quantify the relative abundance of honey bee viruses using the ΔΔCt method. This is because the ΔΔCt method uses honey bee mRNAs for normalization, despite evidence that host RNA degrades more rapidly than viral RNA, often resulting in an over-estimation of virus quantity within postmortem specimens (Dainat et al., 2011; Forsgren et al., 2017). However, many research groups have since moved towards using RT-qPCR with a standard curve method with RNA- or DNA-based tools, to estimate viral genome abundance (Amiri et al., 2019; de Miranda et al., 2021; Dickey et al., 2023; Faurot-Daniels et al., 2020; McCormick et al., 2023; Penn et al., 2021; Runckel et al., 2011). While these methods vary, they all compare virus sequence abundance to an internal or external standard and do not rely on the integrity of the host’s (*i*.*e*., honey bee’s) RNA for accurate quantification. Despite these improved molecular methods to quantify virus abundance, almost all honey bee-associated virus research continues to adamantly follow the “gold-standard” sampling method of collecting live bees because of continued concerns related to RNA degradation. A few studies have demonstrated that postmortem bees can be used to reliably measure virus abundance (Carrillo-Tripp et al., 2016; Dolezal et al., 2016; Hsieh et al., 2020a; Mazzei et al., 2014). However, while these studies found that both live- and postmortem-sampled specimens yielded similar results when measuring virus abundance in honey bees and wild bees, they were not designed to test virus degradation or inactivation, and sampling occurred within a minimal postmortem interval. Collectively, these studies reveal an important gap in our understanding of how prolonged host postmortem intervals affect viral RNA stability, quantification, and continued infectivity.

Developing a better understanding of how virus particles and viral RNA are susceptible to degradation within postmortem bees offers multiple benefits. In addition to providing better insight into questions related to the basic biology of honey bee-associated viruses and their disease dynamics, it would also help researchers make more informed decisions about best sampling practices for both honey bee and wild bee studies. Thus, the ability to quantify virus abundance and determine continued infectivity of honey bee-associated viruses within postmortem hosts requires further investigation. To address this, we tested the hypothesis that Israeli acute paralysis virus (IAPV), a highly virulent and commonly detected honey bee-associated virus (Chen et al., 2014; de Miranda et al., 2010), remains consistently quantifiable and infectious in postmortem honey bees at colony-norms for temperature and humidity. To assess this, we experimentally-infected adult honey bees with IAPV and compared virus abundance in specimens sampled live and at multiple postmortem timepoints. Following this, we performed a second experiment where we extracted virus particles from bees collected at 24- and 48-hours postmortem and tested the infectivity of these extracts via injection into naïve hosts. Overall, our findings demonstrate that IAPV levels are equally quantifiable in honey bees collected live or postmortem, and that IAPV remains viable and infectious within deceased hosts.

## 2. MATERIAL AND METHODS

### 2.1 Virus abundance experiment

To determine if live vs postmortem sampling differentially impacts the accuracy of quantifying virus abundance in honey bees, we experimentally infected bees with IAPV following previously established protocols (Carrillo-Tripp et al., 2016; Hsieh et al., 2020b). In October 2023, honey bees were sourced from colonies managed by the University of Illinois Urbana-Champaign Bee Research Facility in Champaign County, Illinois. Thoracic injections were performed on day-old (<24 hours) adult workers using a purified IAPV inoculum (Hsieh et al., 2020b) that consisted of >99% IAPV purity at a concentration of 2.11 × 10^5^ genome equivalents (g.e.) of IAPV/100 ng of purified total RNA. We chose IAPV as our model virus as aspects of its inoculum potential could be experimentally controlled for, and IAPV is known to cause distinct symptoms within a short infection timeline (Chen et al., 2014; de Miranda et al., 2010). Individual bees were injected with 1 μL of either: 1) the IAPV inoculum diluted to a 10^−6^ dose, or 2) with a heat-killed virus (HKV) aliquot (Carrillo-Tripp et al., 2016) of the same diluted IAPV inoculum to serve as a negative control. Selection of the 10^−6^ dose was chosen based on a previously performed dose-response trial, wherein inoculated bees were still alive within 24 hours post-inoculation (hpi) but exhibited paralysis symptoms indicative of IAPV infection.

Following the injection process, bees were placed into one of two acrylic cages (10 × 10 × 8 cm) based on virus treatment (HKV vs IAPV), with each cage containing 50 injected bees. Both cages contained a gravity sucrose feeder (30% sucrose, w:v) and were kept within an incubator set to in-hive conditions (34 and ∼50% relative humidity) to allow for viral replication to occur. At ∼24 hpi, all bees across both cages were still alive, with all of the IAPV-inoculated bees displaying paralysis symptoms indicative of infection (Chen et al., 2014). At this timepoint, both cages were placed within a -80 ºC freezer for ∼12 hours, halting virus replication and ensuring that each individual across both virus treatments uniformly died at the same time. After this 12-hour period, 10 bees/ virus treatment were kept within the -80 ºC freezer to represent a 0–hour post-death (hpd) sampling timepoint (*i*.*e*., bees sampled live), while the remaining 40 HKV and 40 IAPV bees were removed from -80 ºC conditions, put back into their respective cages, and then placed within the same incubator set to the aforementioned in-hive conditions. Following this, we randomly removed 10 bees from each cage within the incubator at 4, 12, 24, and 48 hpd and then stored them back within -80 ºC conditions until later analysis. In this study, bees were only sampled up to 48 hpd, as previous studies have shown that experimentally-infected honey bees will reach peak IAPV loads at ∼12-36 hpi and then either succumb to or clear infection by 48 hpi (Carrillo-Tripp et al., 2016; Dolezal et al., 2019; Prayugo, 2024). In total, we had 10 treatment groups (n= 2 virus treatments, n= 5 sampling timepoints) with 10 bees/treatment. The only exception was the HKV 48 hpd treatment group, which had one less bee (n= 9) due to low RNA quantity and quality following extraction.

Total RNA was extracted from whole-bodied individuals using the Qiagen RNeasy Mini Kit (QIAGEN) following the manufacturer’s protocol. After extraction, the quantity and quality of total RNA in each sample was measured using a spectrophotometer (Nanodrop 1000). While one bee from the HKV 48 hpd treatment group did not meet the minimum RNA quantity required for our RT-qPCR protocol (≥10 ng of total RNA/μL) and was thus excluded from our abundance analyses, all other samples contained acceptable RNA quantities for RT-qPCR, which included all of the IAPV-inoculated samples. Absolute quantification of IAPV using RT-qPCR was determined via the standard curve method using the primers, standard, and protocol described in Carrillo-Tripp et al. (2016). The only change made here was that we used 20 ng of total RNA as template for each qPCR reaction. The quantity of IAPV in each sample was determined using the QuantStudio RealTime PCR Software (Applied Biosystems, version 1.3, Foster City, CA USA) after completion of the RT-qPCR run. For the statistical analysis of IAPV abundance, IAPV quantities were first log-transformed and then tested for normality using JMP statistical software (JMP®, Pro 17. SAS Institute Inc., Cary, NC, 1989–2023). Significance was determined using the Kruskal-Wallis Test, followed by post-hoc Steel-Dwass tests that protected the overall error rate. The Kruskal-Wallis and Steel-Dwass tests were also performed to determine significance between treatment groups based on mean total RNA concentration (ng/μL). The correlation between sampling timepoint and total RNA concentration within each virus treatment was determined using Spearman’s rank correlation within JMP.

### 2.2 Virus infectivity experiment

To determine whether IAPV remains infectious within honey bees postmortem, we conducted an experiment in May 2024 using similar methods as those described above (section 2.1), including use of the same apiary site, injection protocol, and diluted (10^−6^) HKV and IAPV inoculums. In this experiment, we waited until all of the IAPV-inoculated bees had died naturally as a result of experimental infection to create samples with peak IAPV loads. By 22 hpi, the IAPV cage (n= 20 bees/cage) had reached 100% mortality. At this timepoint (henceforth referred to as <24 hpd due to variation between individuals in their exact time of death), 10 IAPV bees were collected and placed within -80 ºC conditions for later analysis. The remaining 10 IAPV bees were left within the incubator for an additional 24 hours (*i*.*e*. <48 hpd) before being placed within -80 ºC conditions as well. For the HKV control cage, there was 0% mortality by 22 hpi. To create comparable <24 hpd and <48 hpd sampling timepoints that exactly matched the IAPV treatments, the HKV cage was first placed within -80 ºC conditions to uniformly kill all of the HKV bees at 22 hpi. Following this, the resulting postmortem bees were placed back into their respective cage within the incubator and then sampled at the same hpd timepoints as those of the IAPV treatment. In total, we had 4 treatment groups (n= 2 virus treatments, n= 2 sampling timepoints).

From the collected postmortem bees, we created a viral extract for each treatment group using a previously established protocol (Penn et al., 2021; Simone-Finstrom et al., 2018). For each of the 4 treatment groups, 10 whole-bodied bees were homogenized in PBS and then centrifuged at 4,600 rpm at 4 ºC for 30 minutes. The resulting supernatants were then filtered through a 0.22-micron filter to remove tissue debris, fungal spores, and bacteria. With these newly-made viral extracts (*i*.*e*., HKV 24h, HKV 48h, IAPV 24h, and IAPV 48h), we performed a mortality assay using the previously-described injection protocol (section 2.1). This mortality assay also included the source HKV and IAPV inocula (10^−6^) to serve as a negative and positive control, respectively. Each of the six treatment groups consisted of a cage containing injected bees (n=10 bees/cage) that were monitored for mortality over a 48-hour period in 12-hour increments. Pairwise comparisons of survivorship between treatments over the 48 hpi period was performed in RStudio using Cox proportional hazard models with Firth’s Penalized Likelihood corrections (package “coxphf”), followed by Benjamini-Hochberg corrections of the subsequent p-values.

## 3. RESULTS AND DISCUSSION

### 3.1 Virus abundance experiment

We found that IAPV-inoculated bees exhibited significantly higher IAPV loads than our HKV-inoculated controls (Figure 1; Kruskal-Wallis Test; d.f. = 1, χ^2^ = 73.50, *p* < 0.0001). In fact, all of our HKV-inoculated bees had IAPV loads below our RT-qPCR protocol’s limit of detection threshold (Carrillo-Tripp et al., 2016), while all of the IAPV-inoculated bees had mean IAPV loads above what has been previously considered a high-level infection (Harwood et al., 2022). Thus, our controls were putatively free of background IAPV infection, and our experimental injection treatment with IAPV was successful in causing rapid infection. Within the IAPV treatment group, we found that sampling honey bees live (*i*.*e*., 0 hpd) versus at 4, 12, 24, or 48 hpd yielded statistically similar IAPV loads across all pairwise comparisons (Table S1), indicating consistent accuracy in IAPV detection and quantification between live and postmortem bees sampled up to 48 hpd.

**Figure 1:**
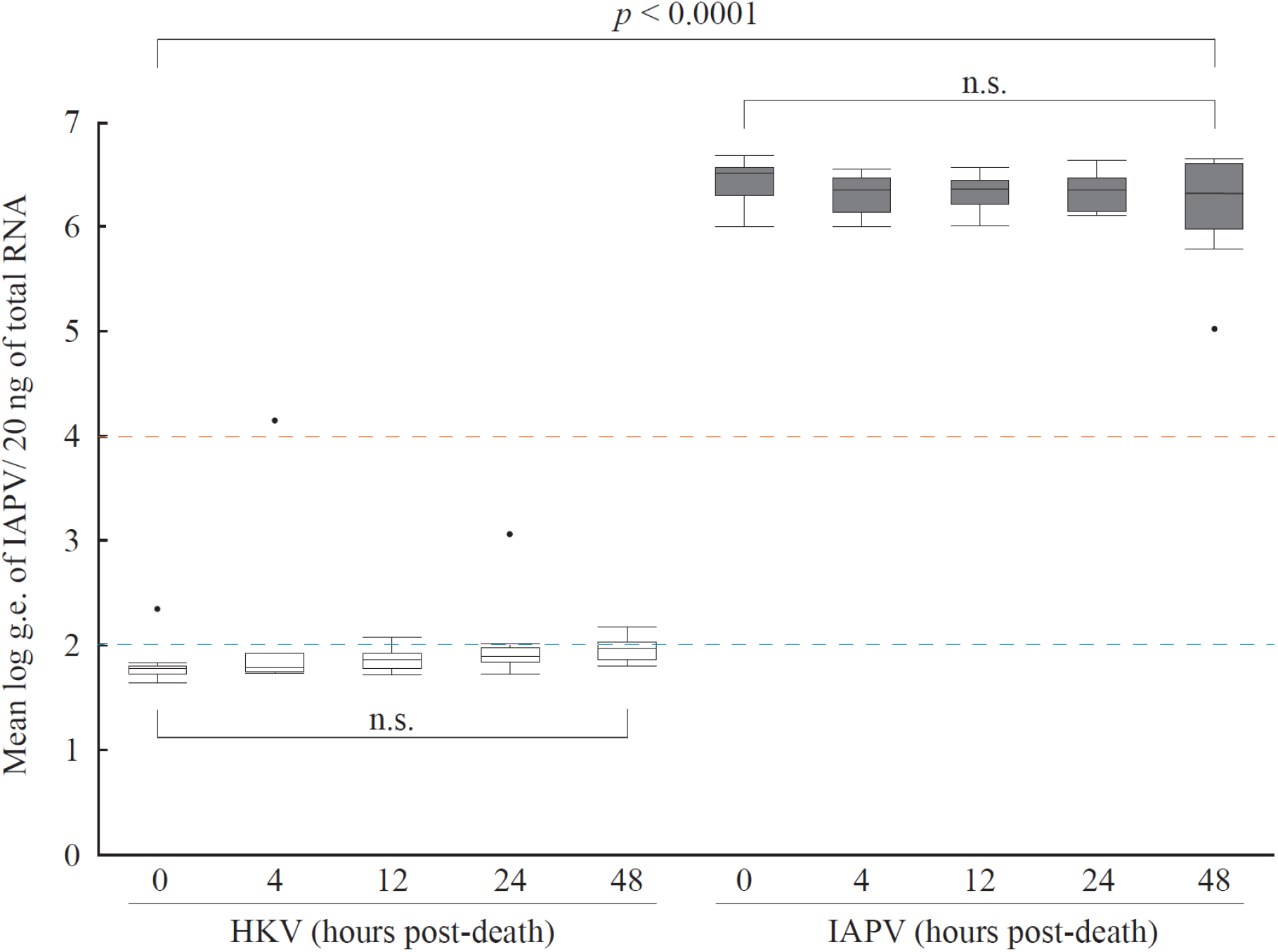
IAPV was detected at similar levels across all sampling timepoints. For each virus treatment (HKV= bees injected with heat-killed IAPV; IAPV= bees injected with infectious IAPV), there were five sampling timepoints (0–, 4–, 12–, 24–, and 48–hours post-death) at which bees were collected to quantify IAPV abundance. The 0–hours post-death (hpd) timepoint represents bees sampled live, while the 4, 12, 24, and 48 hpd timepoints represent bees sampled postmortem. The mean log genome equivalents (g.e.) of IAPV/ 20 ng of total RNA was determined based on individual extractions of whole-bodied adult honey bees (n= 10 bees/treatment group). The HKV 48 hpd treatment group had one fewer bee (n= 9 bees) due to insufficient RNA quantity following extraction. The blue dotted line represents the limit of detection threshold (∼200 g.e. of IAPV/ reaction; Carrillo-Tripp et al., 2016), while the orange dotted line represents the threshold of what has been considered previously as a high-level IAPV infection (>10^4^ g.e./ 20 ng of total RNA; Harwood et al., 2022). Box plots represent the mean (+/- s.e) log g.e. of IAPV and are grouped by virus treatment (HKV= white, IAPV= gray). Significance between the two virus treatments overall (*p* < 0.0001) and between the different sampling timepoints within each virus treatment (n.s.= no significance) are indicated (Kruskal-Wallis test, Steel-Dwass test, α = 0.05).

These results support the use of postmortem bees for quantifying IAPV abundance, which can offer advantages over sampling live bees in certain research scenarios. For instance, virus infections are dynamic and variable, contributing to significant variation and noise in virus loads with random live bee sampling (Carrillo-Tripp et al., 2016; Dolezal et al., 2019; Hsieh et al., 2020b, 2020a). Random sampling of live bees can result in variability when measuring virus abundance due to different response phenotypes, where it is uncertain if any given individual would survive infection. Additionally, focusing solely on live bees can introduce survivor bias (Lipsitch et al., 2015) by overlooking individuals that succumb to infection. Unlike live bees, postmortem bees exhibit a clear phenotype (mortality), likely indicating individual peak virus loads, ensuring greater consistency between sampled individuals. Moreover, sequential sampling of live bees within an experimental group can deplete the population and necessitate censoring in survivorship analysis. Postmortem sampling also provides an alternative in research scenarios where obtaining live specimens is challenging, such as studies involving honey bee-associated viruses in trapped non-*Apis* bees (Dolezal et al., 2016; Fürst et al., 2014; McMahon et al., 2015).

Though IAPV levels were statistically equal between live and postmortem bees, we did observe a negative correlation between sampling timepoint (*i*.*e*., hours post-death) and mean total RNA concentration (ng/μl) within the IAPV virus treatment group (Spearman’s rank correlation; r(48) = -0.43, *p* = 0.0021). While there was a slight significant difference in mean total RNA concentration for the main effect across all five treatment groups (Kruskal-Wallis Test; d.f. = 4, χ^2^ = 9.93, *p* = 0.0416), post-hoc tests (Steel-Dwass test) revealed no significant differences in RNA concentration between any of the sampling timepoints within the IAPV treatment group (Figure 2). While some of the IAPV 48 hpd samples did have relatively low RNA concentrations (∼30 ng/μL), these samples were still sufficient for absolute quantification and resulted in similar levels of IAPV detection compared to samples within the same treatment group that had relatively high concentrations (>1000 ng/μL) of total RNA (Table S2). For the HKV treatment group, we observed a strong negative correlation between sampling timepoint and mean total RNA concentration (Spearman’s rank correlation; r(48) = -0.67, *p* < 0.0001) and significance for the main effect of HKV treatment (Kruskal-Wallis Test; d.f. = 4, χ^2^ = 31.02, *p* < 0.0001). Unlike the IAPV treatment, we also observed significant differences between some of the HKV sampling timepoints after conducting multiple comparison tests (Figure 2).

**Figure 2:**
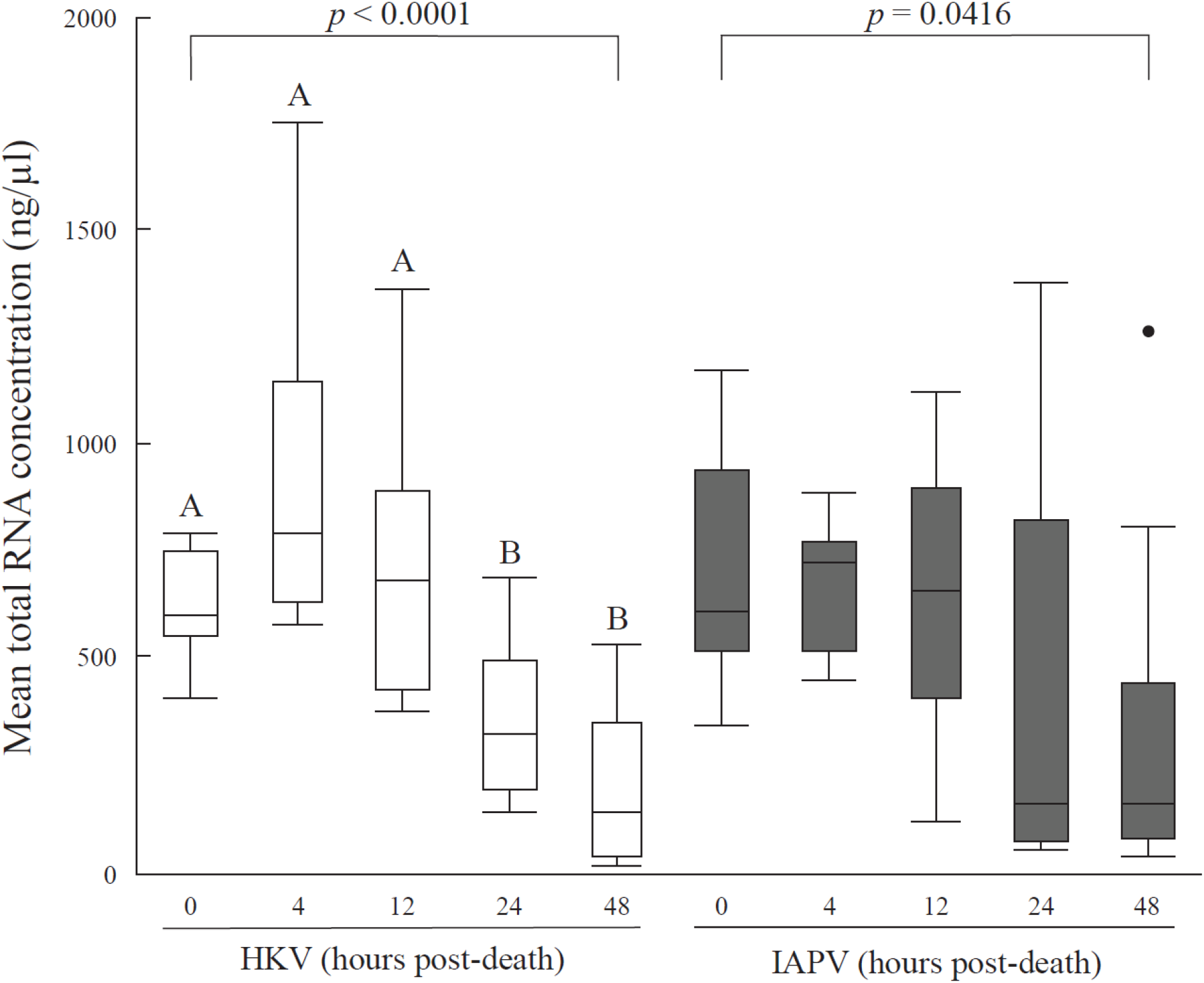
Mean total RNA concentration was similar between the IAPV treatments. The treatment groups consisted of two different virus treatments (HKV= bees injected with heat-killed IAPV; IAPV= bees injected with infectious IAPV) and five sampling timepoints (0–, 4–, 12–, 24–, and 48– hours post-death). For each treatment group, the mean total RNA concentration (ng/μl) was determined based on individual RNA extractions performed on whole-bodied honey bees (n=10 bees/treatment group). This included the HKV 48 hpd bee that fell below the minimum RNA concentration (<10 ng/μl) needed to measure virus abundance with RT-qPCR. Total RNA concentration was determined using a spectrophotometer (NanoDrop 1000). Box plots represent mean (+/-s.e) total RNA and are grouped by virus treatment (HKV= white, IAPV= gray). Significance between the different sampling timepoints within each virus treatment (Steel-Dwass tests; HKV, *p* < 0.0001; IAPV, *p* = 0.0416) are indicated. Different letters within the HKV virus treatment group denote significance within this virus treatment alone. Although there was weak significance overall for the IAPV virus treatment, post-hoc tests showed no significant differences in RNA concentration between any of the sampling timepoints within the IAPV treatment group. For all statistical tests, α = 0.05.

The negative correlations between sampling timepoint and total RNA concentration was expected for both virus treatments, as previous studies have shown that RNA extracted from honey bees degrades over time when not kept at optimal storage conditions (Dainat et al., 2011; Forsgren et al., 2017). However, the consistent quantification of IAPV we observed from postmortem samples, despite evidence of degradation over time of the total RNA, is likely attributed to rapid degradation of host, rather than viral, RNA (Dainat et al., 2011). So, while degradation of the host RNA was occurring, the viral RNA remained intact enough to allow for accurate quantification up to at least 48 hours post-death of the host. Greater degradation of host, and not viral, RNA also helps explain why we found significant differences between the HKV sampling timepoints over time but not between the different IAPV treatment groups. Future studies could better determine this by measuring the extent to which the viral RNA degraded within the postmortem samples, in addition to determining how much time it would take for the postmortem interval to affect the accuracy of quantifying virus abundance.

### 3.2 Virus infectivity experiment

While our first experiment revealed that the quantity of detectable IAPV does not change based on when bees are sampled postmortem, it remained unclear whether these detections were of infectious virus particles, inactivated virus, or simply genome remnants large enough for our primers to detect. To address this, we performed virus particle extractions on <24 and <48 hpd bees and compared their infectivity to our original IAPV inoculum. By injecting each extract into naïve host bees and measuring survivorship over time, we found that IAPV remains highly infectious within postmortem bees for at least 48 hpd (Figure 3). Overall, we found a highly significant effect of virus treatment (HKV vs IAPV) on bee survivorship (Effect Likelihood Ratio Test; d.f. = 1, χ^2^ = 41.34, p < 0.0001). The source HKV inoculum (*i*.*e*., negative control) maintained 100% survival and the HKV 24h and 48h extracts both had 90% bee survival at 48 hpi. Meanwhile, all three of the IAPV treatments (the source IAPV inoculum, IAPV 24h extract, and IAPV 48h extract) had 0% survival by the end of the 48-hour monitoring period, with 0% survival occurring by 12 hpi for both of the IAPV extract treatments and by 24 hpi for the source IAPV inoculum. The significantly higher survivorship observed in the HKV extract treatments compared to the IAPV extracts supports that death of the IAPV-inoculated bees was due to infection. While pairwise comparisons (Table S3) revealed no significant differences between each of the HKV treatments, we did find significant differences in survival between the IAPV treatments, with the source IAPV inoculum having a significantly higher likelihood of survival compared to both the 24h and 48h IAPV extracts (Cox proportional hazards; HR= 11.18, *p* = 0.0008).

**Figure 3:**
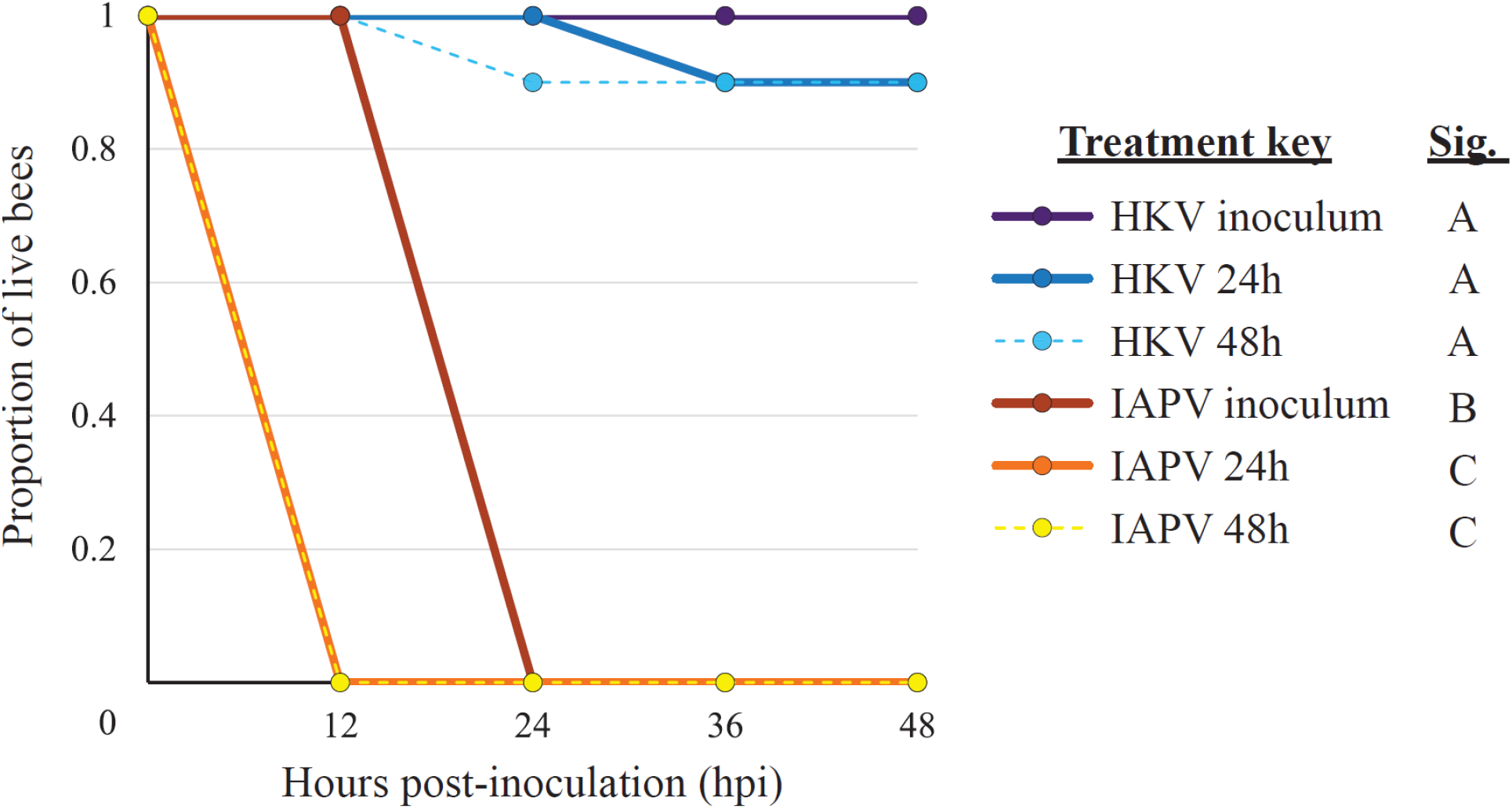
IAPV extracted from postmortem hosts remains highly infectious. Survivorship was measured as the cumulative proportion of live bees/ treatment group determined every 12 hours post-inoculation (hpi) over a 48-hour total monitoring period. Treatment groups were evenly divided between both virus treatments (HKV= bees injected with heat-killed IAPV; IAPV= bees injected with infectious IAPV). Treatments were further divided based on the hour post-death (hpd) at which the postmortem bees were sampled to make the different injection extracts (HKV 24h, HKV 48h, IAPV 24h, and IAPV 48h). The two original inocula that were first used to create the postmortem bees served as control groups (HKV inoculum= negative control, IAPV inoculum= positive control). Each treatment group (n= 6) consisted of single cage of injected day-old bees (n=10 bees/cage). Significance between treatments (Sig.) was determined using pairwise comparisons of Cox proportional hazard models (α = 0.05) and is depicted by differing letters adjacent to the treatment key.

These results show that some proportion of IAPV particles remain infectious in postmortem hosts for at least 48 hpd. While we cannot discern what the exact proportion of infectious particles is, the IAPV extracts causing greater mortality risk than the source IAPV inoculum suggests that significantly high quantities of IAPV still remain infectious post-host death. This supports the hypothesis that IAPV remains fairly resistant to degradation, even within a deceased host kept at warm and humid conditions (34 and ∼50% relative humidity), at least for 48 hours post-death. Our results suggests that honey bees that have died as a result of IAPV infection can continue to act as a source of transmission for healthy individuals within a colony and underlines the importance of the evolution of social immune behaviors aimed at reducing corpse-mediated transmission, such as corpse-removal behavior (López-Riquelme and Fanjul-Moles, 2013) performed by specialized undertaker honey bees within a colony (Trumbo et al., 1997; Visscher, 1983). It also supports corpse-mediated transmission as a viable route through which parasites and predators of bee species may acquire infectious honey bee-associated viruses (Payne et al., 2020; Yañez et al., 2020, 2012; Yang et al., 2020).

## 4. CONCLUSION

Results from this study showed that IAPV remains highly infectious within postmortem honey bees for up to 48 hours post-death, supporting the hypothesis that the quantifiable IAPV in postmortem hosts contains active virus rather than only inactivated virus or genome remnants. This continued infectiousness of IAPV implies that it has considerable resistance to degradation within postmortem hosts, even under hot and humid conditions resembling honey bee colony environments. Our findings also supported the hypothesis that postmortem honey bee sampling yields comparable results to live bee sampling when quantifying IAPV abundance using the standard curve method. In fact, postmortem sampling offers several advantages over live sampling when quantifying viral load, including minimizing variability between samples, mitigating survivor bias, preserving the overall sample pool, and enabling research questions where only postmortem specimens are available. As the scope of this study focused on IAPV within its typical 48-hour infection timeline (Carrillo-Tripp et al., 2016; Dolezal et al., 2019; Prayugo, 2024) further studies will be needed to determine the full extent to which IAPV remains infectious within a postmortem host, in addition to whether these same conclusions can apply to other bee viruses and host species. Overall, these findings support the use of postmortem bees for quantifying IAPV abundance and highlight the continued infectiousness of IAPV within postmortem hosts, suggesting corpse-mediated transmission as a probable route for IAPV persistence and infection of naïve hosts.

## ACKNOWLEDGEMENTS

We would like to thank Rachel Manweiler, Sarah M. Murphree, and Nathanael J. Beach for honey bee colony management, and we would like to thank Edward M. Hsieh for providing a graphic design of the cages used in this experiment.

## AUTHOR CONTRIBUTIONS

Conceptualization: AGD; Data curation: ANP, VP; Formal analysis: ANP, VP; Funding acquisition: AGD; Investigation: ANP, VP; Methodology: ANP, VP; Project administration: ANP; Resources: AGD; Supervision: AGD; Visualization: ANP; Writing - original draft: ANP; and Writing - review & editing: ANP, VP, AGD.

## FIGURE LEGENDS

**Table S1:** All pairwise comparisons of quantified IAPV abundance between treatment groups for the virus abundance experiment using post-hoc Steel-Dwass tests.

**Table S2:** Ct values and IAPV quantity of each individual bee after completion of RT-qPCR for the virus abundance experiment.

**Table S3:** All pairwise comparisons determining significance in survivorship between treatments in the virus infectivity experiment. Significance was determined using mixed effect Cox proportional hazard models followed by Benjamini-Hochberg corrections. Asterisks denote significant Level 1/Level 2 pairwise comparisons.

## GRAPHICAL ABSTRACT

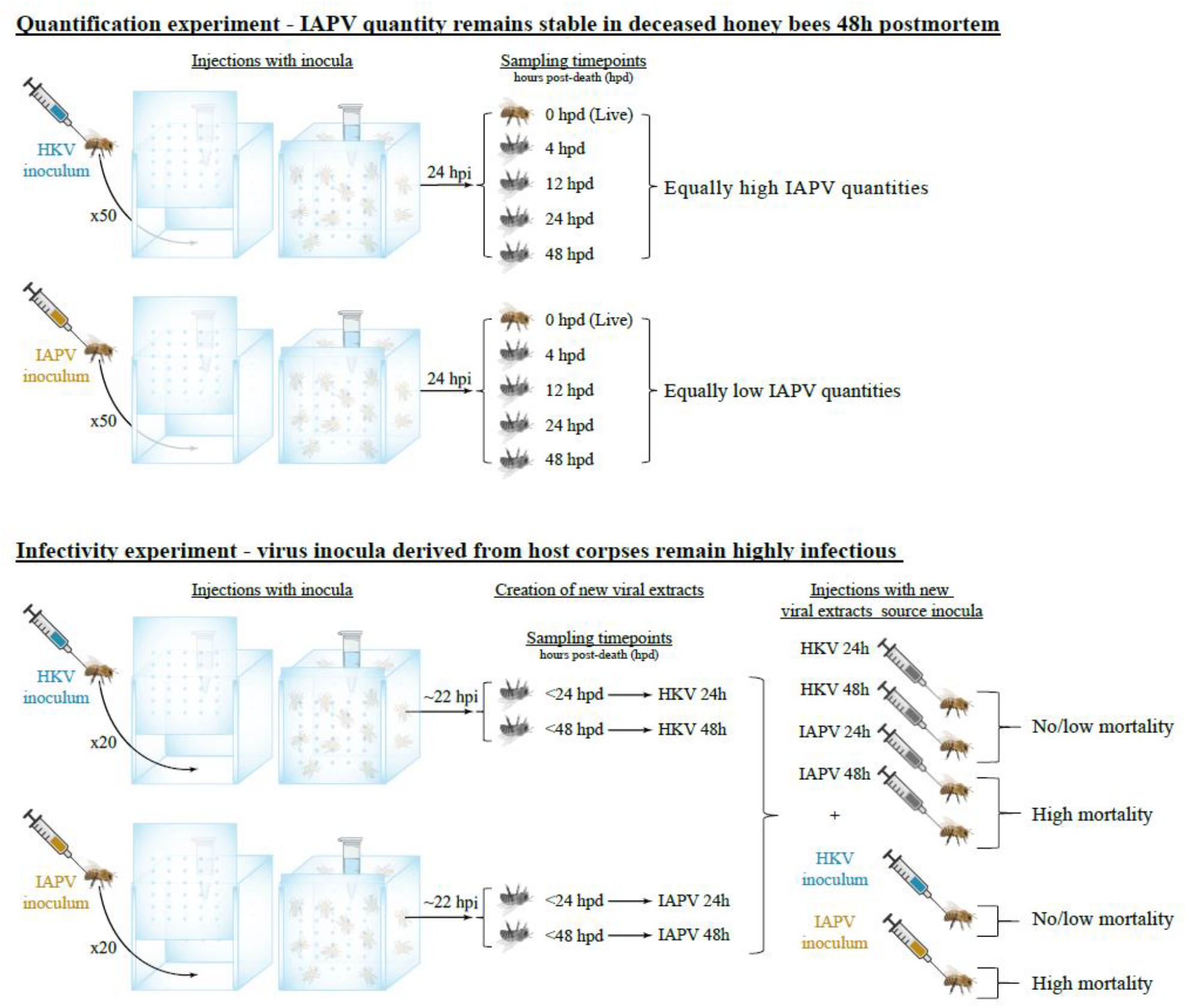

